# Molecular basis for the evolution of species-specific hemoglobin capture by pathogenic *Staphylococcus*

**DOI:** 10.1101/339705

**Authors:** Jacob E. Choby, Hanna B. Buechi, Allison J. Farrand, Eric P. Skaar, Matthew F. Barber

**Affiliations:** Department of Pathology, Microbiology, and Immunology, Vanderbilt University Medical Center, Nashville, TN, 37232,USA; Vanderbilt Institute for Infection, Immunology, and Inflammation, Vanderbilt University Medical Center, Nashville, TN, 37232, USA; Graduate Program in Microbiology and Immunology, Vanderbilt University, Nashville, TN, 37232, USA; Institute of Ecology and Evolution, University of Oregon, Eugene, OR, 97403, USA

## Abstract

Metals are a limiting resource for pathogenic bacteria and must be scavenged from host proteins. Hemoglobin provides the most abundant source of iron in the human body and is required by several pathogens to cause invasive disease. However, the consequences of hemoglobin evolution for bacterial nutrient acquisition remain unclear. Here we show that the α- and β-globin genes exhibit strikingly parallel signatures of adaptive evolution across simian primates. Rapidly evolving sites in hemoglobin correspond to binding interfaces of IsdB, a bacterial hemoglobin receptor encoded by pathogenic *Staphylococcus aureus*. Using an evolution-guided experimental approach, we demonstrate that divergence between primates and staphylococcal isolates governs hemoglobin recognition and bacterial growth. Reintroducing putative adaptive mutations in α- or β-globin proteins is sufficient to impair *S. aureus* binding, providing a mechanism for the evolution of disease resistance. These findings suggest that bacterial hemoprotein capture has driven repeated evolutionary conflicts with hemoglobin during primate descent.

## INTRODUCTION

Animals possess a variety of molecular factors that effectively sequester essential metals from invasive microbes, contributing to an innate immune function termed nutritional immunity (1, 2). Iron, as a critical cofactor for many host and bacterial enzymes, has provided the paradigm for our current understanding of nutritional immunity. Since the discovery of iron limitation by egg white ovotransferrin in the 1940s (3), mechanisms underlying nutritional immunity and bacterial iron scavenging have been the subject of intense study (4). Many vertebrate-associated bacteria encode high affinity uptake systems targeting heme, an abundant iron-containing porphyrin cofactor (5).

The most abundant source of heme-iron in the mammalian host is hemoglobin, which mediates oxygen transport within circulating erythrocytes. The predominant adult hemoglobin consists of a tetramer containing two α-globin and two β-globin protein subunits, each of which binds a single heme molecule for coordination of oxygen. The Gram-positive bacterium *Staphylococcus aureus* is well-adapted to the human host and is a leading cause of skin and soft tissue infections, endocarditis, osteomyelitis, and bacteremia (6). In order to acquire iron during infection, *S. aureus* has evolved a high-affinity hemoglobin binding and heme extraction system, termed the *i*ron regulated surface determinant (Isd) system (7). Following the lysis of proximal erythrocytes via secreted bacterial toxins, released hemoglobin is captured by receptors at the *S. aureus* cell surface (8, 9). The Isd system of *S. aureus* in part consists of cell-wall anchored IsdB and IsdH, which bind hemoglobin and haptoglobin-hemoglobin, respectively (9, 10).

We and others have shown that IsdB is the primary hemoglobin receptor for *S. aureus* and critical for pathogenesis in murine models (9, 11–13). Additionally, IsdB is highly expressed in human blood (14) and a promising vaccine target (15), underscoring its importance in human disease. IsdB extracts heme from hemoglobin, and heme is subsequently passed across the cell wall and into the cytoplasm for degradation by the heme oxygenases IsdG and IsdI, liberating iron (16–20). Underscoring the importance of IsdB for pathogenesis, heme is the preferred iron source of *S. aureus* during murine infection (21). The cell-wall anchored IsdABCH proteins share between one and three NEAT (*nea*r transporter) domains for coordination of hemoglobin or heme. IsdB NEAT1 binds hemoglobin while NEAT2 binds heme, tethered by an intervening linker (22). Consistent with adaptation of *S. aureus* to colonize and infect humans, we previously found that *S. aureus* IsdB binds human hemoglobin more effectively than mouse hemoglobin, the common laboratory animal used to model *S. aureus* infection (13). These results suggest that hemoglobin variation among mammals could dictate effective heme acquisition by *S. aureus* and other Gram-positive bacteria.

Previous work has demonstrated that pathogens can promote rapid adaptation of host immunity genes through repeated bouts of positive selection (23–25). While adaptation during such evolutionary conflicts can take many forms, theoretical and empirical studies concur that an elevated rate of nonsynonymous to synonymous substitutions in protein-coding genes is indicative of recurrent positive selection (26, 27). To date, most empirical studies of host-pathogen ‘arms races’ have focused on viruses (28–31). Recently we showed that the transferrin family of iron-binding proteins has undergone extremely rapid evolution in primates at protein surfaces bound by iron-acquisition receptors from Gram-negative bacteria (32, 33). These findings are consistent with a long-standing evolutionary conflict for nutrient iron, whereby mutations in iron binding proteins that prevent bacterial scavenging protect the host from infection and are favored by natural selection. While these studies have expanded our understanding for how pathogens shape the evolution of host genomes, they also raise the question of whether other components of nutritional immunity might be subject to similar evolutionary dynamics.

In addition to its role as the principal bloodstream oxygen transporter, hemoglobin has provided an important biological model for diverse areas of the life sciences. Elegant studies have illustrated how hemoglobin variation underlies multiple instances of adaptation to high altitudes in diverse vertebrate taxa (34–37). Hemoglobin alleles have also likely been subject to balancing selection in human populations, where mutations that produce sickle-cell disease also confer resistance to severe malaria (38) and have reached high frequencies in regions where malaria is endemic. Despite its long history of study, the consequences of hemoglobin evolution for vertebrate nutritional immunity remain unclear. In the present study we set out to investigate the evolution of hemoglobin family proteins in primates and determine whether primate hemoglobin evolution impacts the ability to sequester heme-iron from bacterial pathogens.

## RESULTS

### Parallel signatures of positive selection in primate hemoglobins at the IsdB binding interface

To investigate how natural selection has shaped hemoglobin diversity in simian primates, orthologs of the α- and β-globin genes were cloned and Sanger sequenced from primate cell lines as well as compiled from publically-available databases. In total, 27 α-globin and 30 β-globin orthologs were assembled for phylogenetic analyses using the PAML and HyPhy software packages (Figure 1A, Materials and Methods), which use nonsynonymous and synonymous substitution rates to infer signatures of positive selection. Because globin genes have been shown to undergo gene conversion which can distort inferred phylogenetic relationships, all analyses were performed using both a well-supported species tree as well as gene trees generated using PhyML. All tests detected significant evidence of positive selection acting on both α- and β-globin using both species and gene phylogenies (Supplemental Data). Multiple iterations repeatedly identified two sites in α- and β-globin exhibiting strong signatures of positive selection (Figure 1A). It was immediately apparent that these rapidly-evolving sites localized to similar regions of the α- and β-globin proteins, specifically the N-terminal A helix and the hinge region between the E and F helices (Figure 1B). In fact, the two sites exhibiting signatures of selection in the α- and β-globin A helices are at homologous positions. These parallel signatures of selection between α- and β-globin could indicate that a similar selective pressure has driven this divergence between primate species. To investigate whether bacterial heme scavenging receptors could be one such pressure, rapidly evolving sites were mapped onto a recently solved co-crystal structure between human hemoglobin and the IsdB protein from *S. aureus*. Notably, all four residues are localized to the IsdB binding interface, in close proximity to the NEAT1 domain (Figure 1C). Together these findings indicate that primate globins have undergone rapid divergence at specific sites proximal to the binding interface with Gram-positive bacterial hemoglobin receptors.

**Figure 1.**
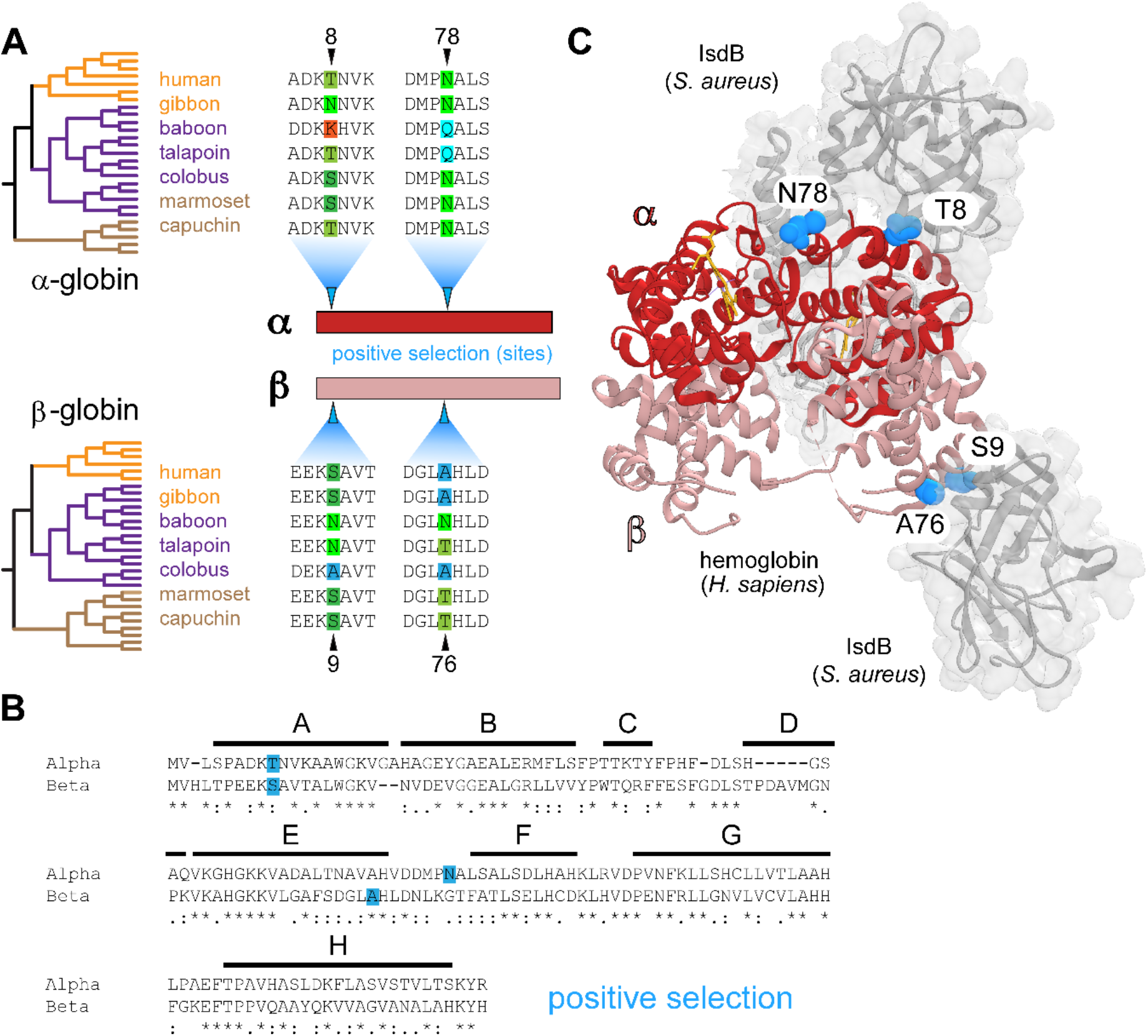
Parallel signatures of positive selection among primate hemoglobins at the bacterial IsdB binding interface. (A) Species phylogenies for α-globin and β-globin orthologs analyzed (left) with alignments from representative species across hominoid (orange), Old World monkey (purple) and New World monkeys (brown). Amino acid sites under positive selection were identified in α-globin and β-globin (blue arrows) using PAML with both species and gene trees (posterior probabilities >0.95). (B) Amino acid alignment of human α-globin and β-globin proteins. The position of conserved helices A-H are shown, and identity between globins is noted as conserved (*), highly similar (:), and similar (.). (C) The residues of α-globin (red) and β-globin (salmon) under positive selection (blue spheres) at the interface of hemoglobin capture by *Staphylococcus aureus* IsdB (gray) (PDB:5VMM).

### Primate hemoglobin variation dictates *S. aureus* binding and heme-iron acquisition

To assess how hemoglobin divergence among primates impacts recognition by *S. aureus*, recombinant hemoglobin from human, white-cheeked gibbon, baboon, talapoin, and marmoset were purified, providing broad representation from our phylogenetic dataset. An established biochemical assay was used to measure binding of hemoglobin by *S. aureus*, in which *S. aureus* cells recognize recombinant human hemoglobin as well as hemoglobin purified from blood in an IsdB-dependent manner (Supplemental Figure 1). *S. aureus* exhibited significantly reduced binding of baboon and marmoset hemoglobin to the cell surface (Figure 2A). It was noted that binding patterns do not strictly match predictions based on host phylogeny, suggesting discrete large-effect substitutions in hemoglobin may contribute disproportionately to recognition by *S. aureus*. We next determined the ability of primate hemoglobins to support growth of *S. aureus* as the sole iron source. Consistent with whole-cell binding data, hemoglobins that were bound by *S. aureus* with low affinity were unable to support optimal bacterial growth, indicating that the capability to bind hemoglobin is a measure of the ability to utilize hemoglobin as an iron source (Figure 2B). Together these results demonstrate that variation among primate globins dictates bacterial hemoglobin capture and heme-dependent growth.

**Figure 2.**
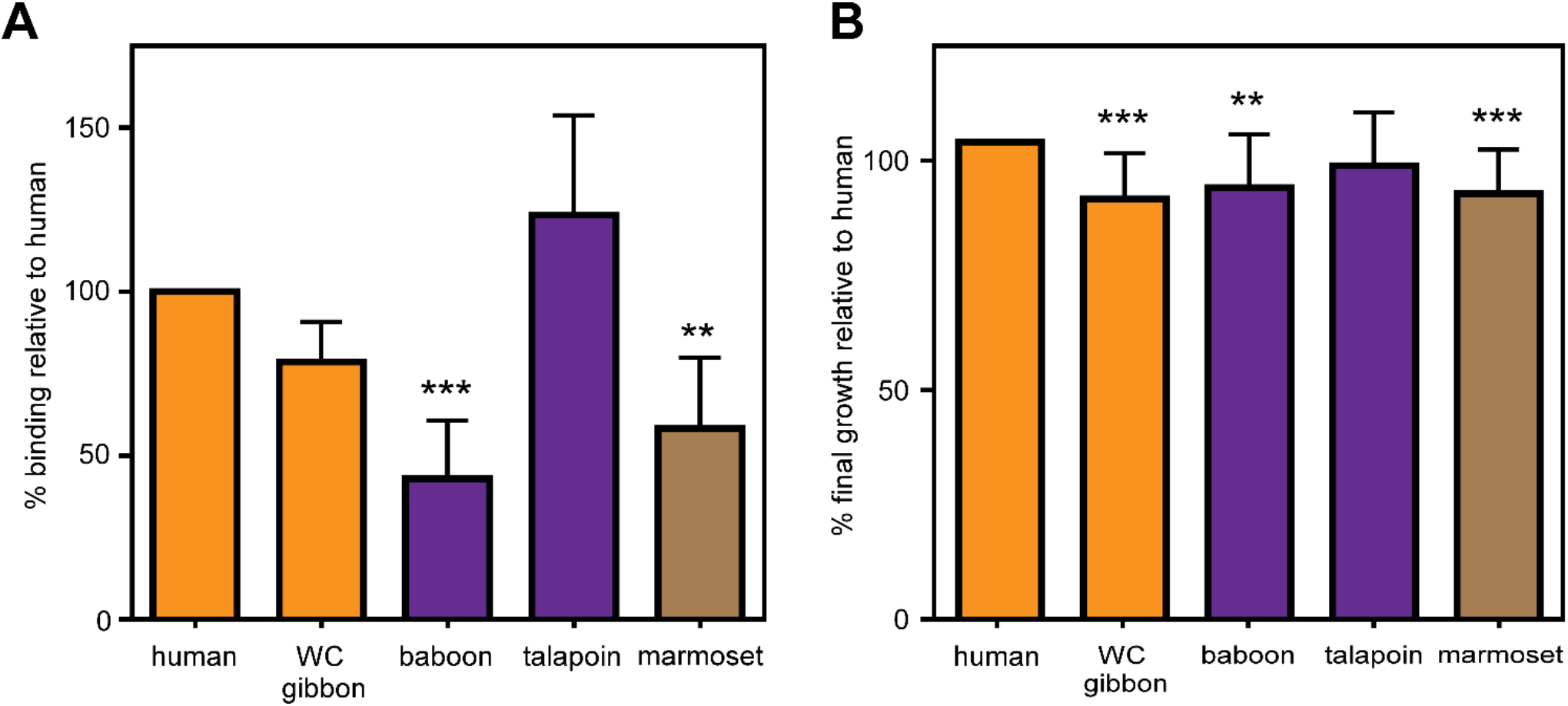
Primate hemoglobin variation dictates *S. aureus* binding and heme-iron acquisition. (A) *S. aureus* binds hemoglobin with species specificity. Iron-starved *S. aureus* was incubated with 10 μg/ml purified recombinant hemoglobin from representative species. Hemoglobin bound to the surface of *S. aureus* was eluted and analyzed by SDS-PAGE; relative hemoglobin abundance was measured by densitometry analysis (Image J) and compared to human hemoglobin for each replicate. (B) IsdB preference of hemoglobin binding correlates with the ability of *S. aureus* to utilize hemoglobin as the sole iron source. Growth of *S. aureus* in iron-deplete medium with 2.5 μg/ml of purified recombinant hemoglobin as the sole iron source. Shown is the final growth yield of *S. aureus* after 48 hours. For A and B, graphed are the means of four independent experiments in biological triplicate +/− SEM, ** *p*<0.005; *** *p*<0.0005 by two-way ANOVA with Sidak correction for multiple comparisons.

### Species-specific diversity in α-globin restricts heme scavenging by *S. aureus*

The identification of rapidly evolving sites at the IsdB binding interface in both α-globin and β-globin suggest that natural selection in primate globins has been driven by antagonistic evolutionary conflicts with related families of bacterial receptors during primate divergence. Mechanistically, these findings also suggest that both globins contribute to *S. aureus* species-specific hemoglobin capture. We therefore exploited the enhanced binding of human hemoglobin relative to baboon to examine the role of each globin subunit in this biochemical interaction. The ability of *S. aureus* to bind chimeric hemoglobins was measured, which revealed that both globins contribute to species-specificity (Figure 3A), as chimeras containing either human α- or β-globin were bound more effectively than baboon hemoglobin. However, α-globin appears to have a greater effect on human-specific capture, as the α-human β-baboon chimera was bound significantly better than α-baboon β-human. Focusing on phylogenetic variation at the protein binding interface, α-globin T8 and N78 are both proximal to the NEAT1 domain of IsdB (Figure 3B, 3C). Mutagenesis of the N-terminal alpha helix of human α-globin revealed that substituting the Lys of baboon at position 8 significantly reduced binding by *S. aureus* (Figure 3D). Additionally, substituting A5D, T8K, and N9H in human α-globin, which converts this 7 amino acid region to that of baboon, leaves *S. aureus* binding indistinguishable from that of baboon hemoglobin. These results demonstrate that the N-terminal helix of α-globin makes a major contribution to *S. aureus* hemoglobin specificity. Next, the relative importance of the rapidly evolving N78 residue in α-globin was assessed, which lies N-terminal to the sixth alpha helix (Figure 3B). Substitution of N78 to glutamine (present in baboon, talapoin, and other Old World primates) or to histidine, reduced binding of human hemoglobin (Figure 3E). Thus, substitutions at multiple residues in α-globin that exhibit signatures of repeated positive selection disrupt the ability of *S. aureus* to recognize human hemoglobin.

**Figure 3.**
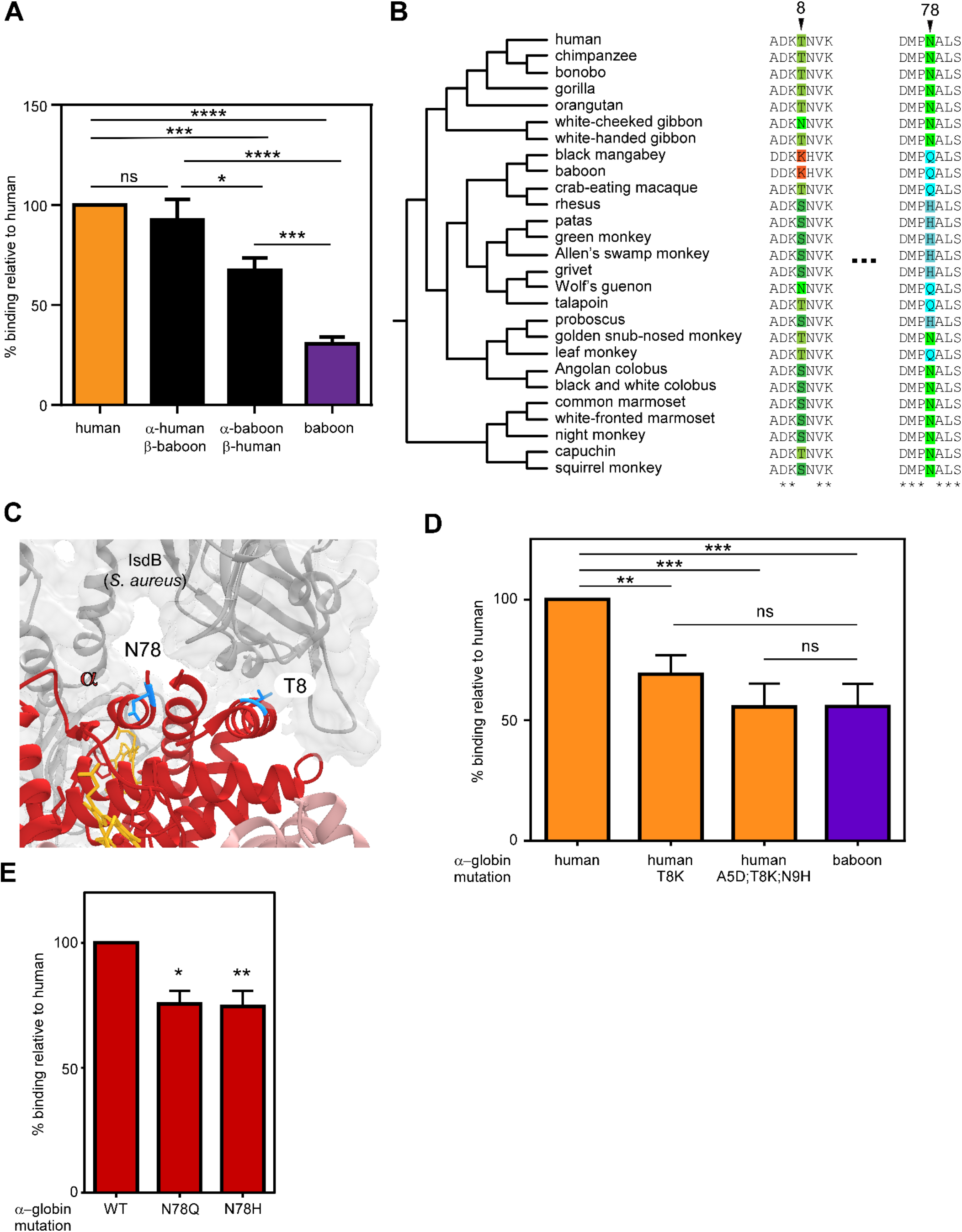
Species-specific diversity in α-globin restricts heme scavenging by *S. aureus*. (A) *S. aureus* binding of human, baboon, and chimeric hemoglobins demonstrates that both globins contribute to binding by IsdB. (B) Species phylogenies and sequence alignments surrounding positions exhibiting signatures of positive selection in α-globin. (C) Residues 8 and 78 of human α-globin (red) interact closely with IsdB (gray) (PDB:5VMM). (D) The T8 residue in α-globin contributes to IsdB recognition between human and baboon hemoglobin as assessed by IsdB binding of mutagenized hemoglobin. (E) The N78 residue of human α-globin contributes to species specificity by IsdB. For A, D, E, graphed are the means of 3-4 independent experiments with 2-3 biological replicates +/− SEM, ns: no significance, * *p*<0.05, ** *p*<0.005; *** *p*<0.0005 by two-way ANOVA with Sidak correction for multiple comparisons.

### β-globin divergence contributes to *S. aureus* hemoglobin binding

*S. aureus* was capable of binding α-baboon β-human chimeric hemoglobin with higher affinity than baboon hemoglobin (Figure 3A), signifying that β-globin also contributes to *S. aureus* species-specific hemoglobin capture. Therefore, the contribution of rapidly evolving residues in β-globin to this binding interaction were investigated (Figure 4A). Both S9 and A76 interact closely with the NEAT1 domain of IsdB (Figure 4B). The effect of substituting human β-globin S9 and A76 with residues found in other primate species analyzed in this work was systematically tested, which revealed that A76 is particularly important for binding by *S. aureus* (Figure 4C). Notably, baboon hemoglobin displays phylogenetic variation at both position 9 and 76, suggesting that these residues may contribute to the inability of IsdB to bind baboon hemoglobin. This variance might also explain the binding affinity differences between human hemoglobin and the α-human β-baboon chimera, observed in Figure 3A. As for α-globin, no single residue substitution improved binding by *S. aureus* IsdB, consistent with the idea that IsdB has specifically adapted to bind human hemoglobin. Taken together with earlier data, residues at the IsdB interface of both α-globin and β-globin contribute to recognition of hemoglobin by *S. aureus*. This is consistent with the NEAT1 domain of multiple IsdB monomers engaging in hemoglobin capture by binding both α- and β-globins, as observed in the co-crystal structure (22).

**Figure 4.**
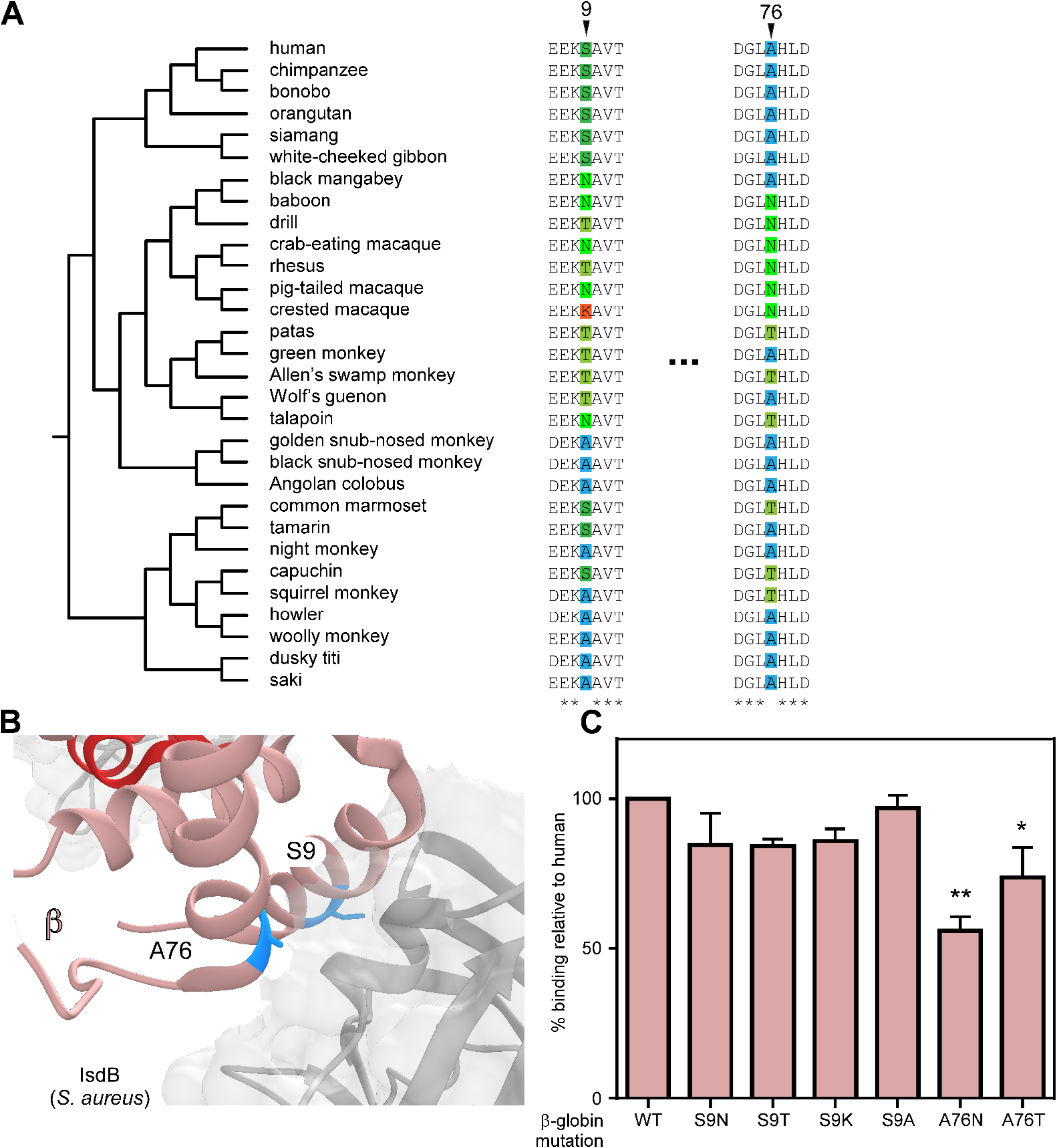
β-globin divergence contributes to *S. aureus* hemoglobin binding. (A) Species phylogenies and sequence alignments surrounding positions exhibiting signatures of positive selection in β-globin. (B) Residues 9 and 76 of human β-globin (salmon) interact closely with IsdB (gray) (PDB:5VMM). (C) The identity of the positively selected residues in human β-globin impact binding by *S. aureus;* β-globin residues S9 and A76 were mutagenized in human hemoglobin. Graphed are the means of four independent experiments with 2-3 biological replicates +/− SEM, * *p*<0.05; ** *p*<0.005 by two-way ANOVA with Sidak correction for multiple comparisons.

### IsdB diversity among related staphylococcal strains impacts primate-specific hemoglobin capture

Given the observed differences in *S. aureus* binding between diverse primate hemoglobins, we considered how genetic variation in IsdB might impact this interaction. The IsdB NEAT1 subdomain Q162R-S170T is critical for hemoglobin recognition and is completely conserved among more than three thousand *S. aureus* clinical isolates (11). Therefore, IsdB variation among congeneric *S. argenteus* and *S. schweitzeri* was assessed. These recently diverged taxa (Figure 5A) are both primate-associated and, unlike most other staphylococci, encode IsdB. We measured the ability of IsdB from *S. argenteus* and *S. schweitzeri* to bind hemoglobin by expressing them ectopically in *S. aureus*. Consistent with their overall high sequence identity, *S. schweitzeri* and *S. argenteus* IsdB bind primate hemoglobin with a similar pattern of species preference as *S. aureus* (Figure 5B). However, both the IsdB of *S. schweitzeri* and *S. argenteus* display reduced binding of talapoin hemoglobin, and *S. argenteus* IsdB does not bind marmoset significantly worse than human. These data indicate that variation among IsdB impacts species-specific hemoglobin capture.

**Figure 5.**
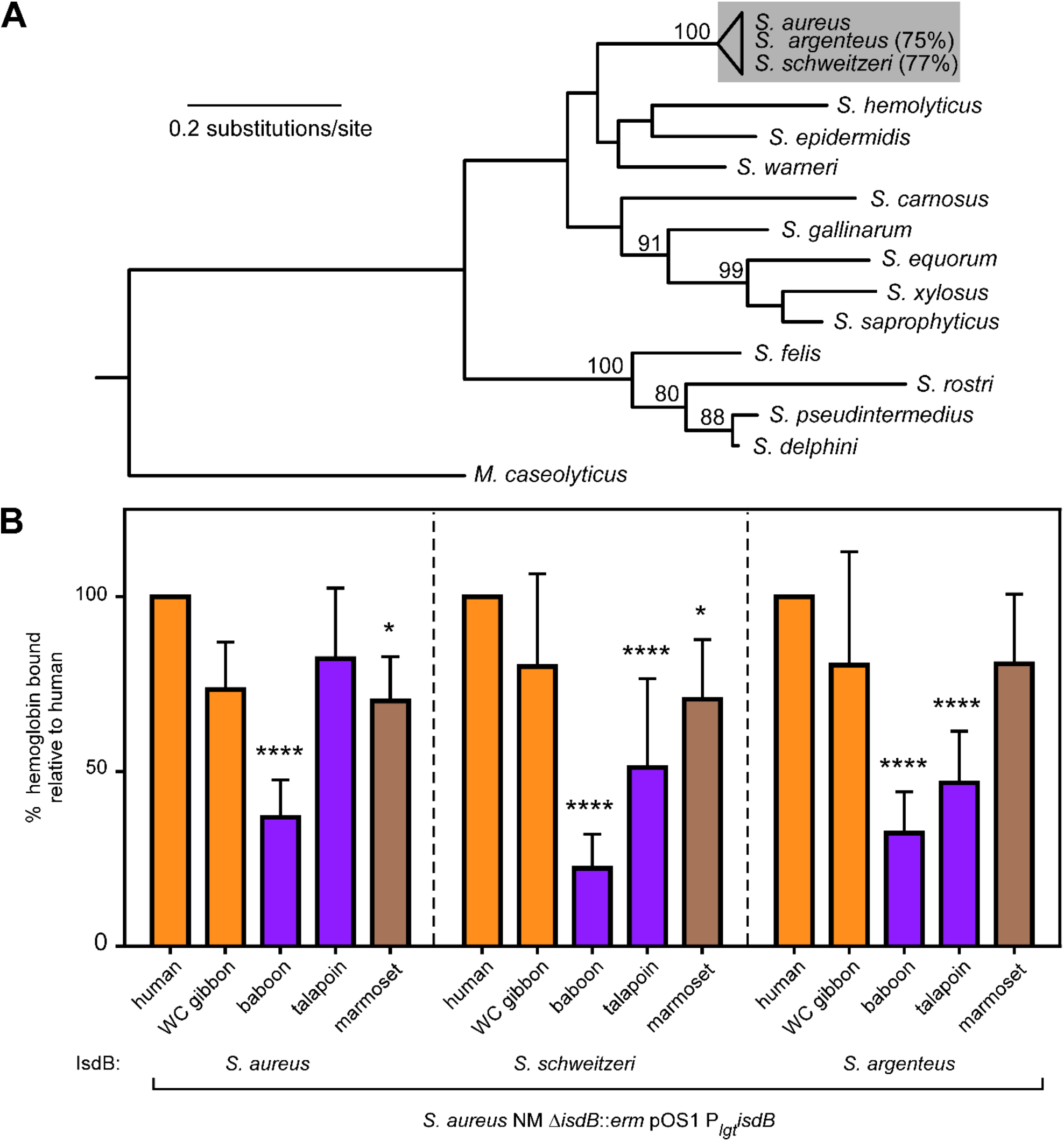
IsdB diversity among related staphylococcal strains impacts primate-specific hemoglobin capture. (A) Maximum likelihood phylogeny of the DNA gyrase A protein from representative staphylococci generated using PhyML. *M. caseolyticus* was included as an outgroup. The similarity of IsdB in *S. argenteus* and *S. schweitzeri* relative to *S. aureus* is shown at right. Bootstrap values above 80 are indicated. B) The IsdB of *S. aureus, S. schweitzeri*, and *S. argenteus* demonstrate a common species preference for hemoglobin binding when expressed ectopically in *S. aureus*. Graphed are the means of three independent experiments with 3 biological replicates +/− SEM, * *p*<0.05; **** *p*<0.0001 by two-way ANOVA with Sidak correction for multiple comparisons.

### IsdB NEAT1 domain diversity among staphylococci modulates human hemoglobin recognition

Closer examination of the Q162R-S170T region of IsdB NEAT1 revealed variation between related staphylococci, but no variation in the critical heme-binding region of NEAT2 (Figure 6A). This region of NEAT1 closely interacts with the N-terminal helices of either α-globin and β-globin, in close proximity to both discrete sites bearing signatures of adaptive evolution in α-globin and β-globin (Figure 6B). To determine the functional consequences of variation in this NEAT1 domain, Q162 and S170T were mutagenized in *S. aureus* IsdB to mimic the sequence of *S. schweitzeri* and *S. argenteus*. Variations at both of these positions reduced affinity for human hemoglobin, showing that in the context of *S. aureus* IsdB, Q162 and S170 are required for high affinity hemoglobin binding (Figure 6C).

**Figure 6.**
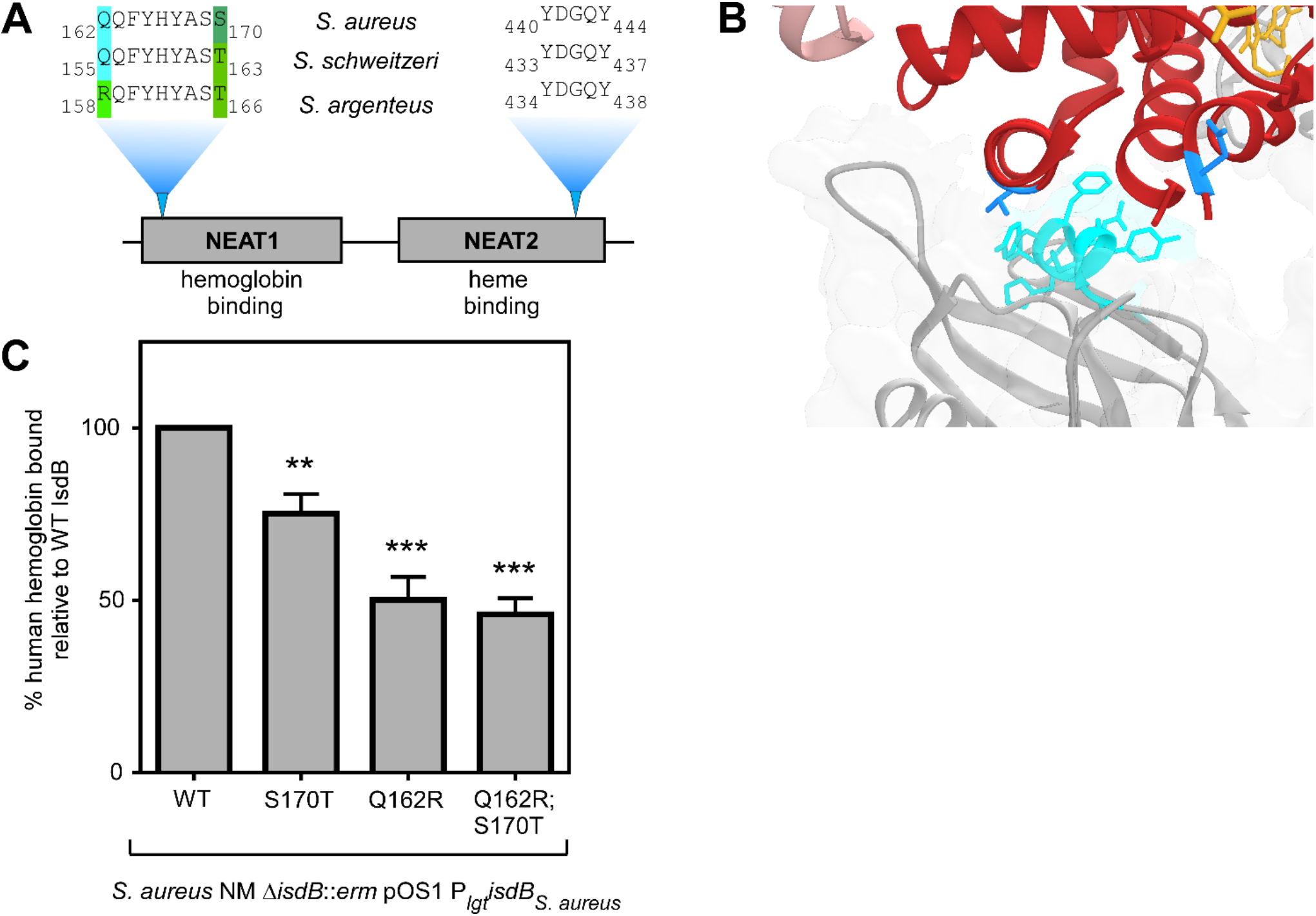
IsdB NEAT1 domain diversity among staphylococci modulates human hemoglobin recognition. (A) An alignment of the NEAT1 subdomain critical for hemoglobin binding shows variation among staphylococcal IsdB, while no variation is observed for the NEAT2 subdomain required for heme binding. (B) The Q162-S170 subdomain of NEAT1 (cyan) is proximal to helices containing T8 and N78 of α-globin (red). (C) The NEAT1 subdomain of *S. aureus* is required for high affinity human hemoglobin binding. Plasmid encoded IsdB was mutagenized and the impact of variation at the terminal residues of the NEAT1 subdomain was assessed by hemoglobin binding. Graphed are the means of three independent experiments with 3 biological replicates +/− SEM, ** *p*<0.005, *** *p*<0.0005 by two-way ANOVA with Sidak correction for multiple comparisons.

## DISCUSSION

In this work we report that recurrent positive selection acting on primate α- and β-globin proteins restricts hemoglobin binding and nutrient acquisition by pathogenic *S. aureus*. Estimations of divergence in the *Staphylococcus* genus have been lacking, however the Kloos hypothesis (39) contends that staphylococci have coevolved with their mammalian hosts over long evolutionary timescales. In support of this concept, primate-specificity among staphylococci has been reported including *S. aureus*, *S. epidermidis*, and *S. warneri*, as well as avian (*S. gallinarum*), equine (*S. equorum*), and others. Indeed, it has been proposed that the canine-associated *S. pseudointermedius* diverged from *S. aureus* simultaneously with the divergence of Primate and Carnivora orders (40). Most staphylococci are commensal organisms, while *S. aureus* is uniquely adapted to infect deep tissue and cause disease. As such, the IsdB system is only encoded by *S. aureus* and closely related primate-associated staphylococci. By narrowing our analysis of hemoglobin evolution to primates, we are thus able to assess specific biological features of primate-associated staphylococci.

An outstanding question in the study of *S. aureus* evolution has been determining the selective pressures responsible for human-specific virulence factors. *S. aureus* asymptomatically colonizes the anterior nares of approximately one third of the human population, yet is capable of causing a wide range of invasive diseases. While some bacterial colonization factors have been implicated in pathogenesis, many virulence factors have evolved highly specific targets that are not obviously involved in nasal colonization (41). Additionally, transmission has been proposed to be a selective pressure that enhanced the virulence of *S. aureus* relative to another human skin commensal, *S. epidermidis* (42). As such, we cannot definitively conclude that IsdB evolution has been driven by selection during invasive disease. It is also likely that variation across IsdB of *S. aureus, S. argenteus*, and *S. schweitzeri* may be the results of antigenic variation to evade the immune system. By focusing on the hemoglobin binding pocket of IsdB, we have been able to pinpoint critical variation for hemoglobin specificity.

Our phylogenetic analyses revealed strikingly parallel signatures of positive selection between the α- and β-globin genes across primates. In particular, rapidly evolving sites in the α- and β-globin A-helices are predicted to be homologous based on predicted protein alignments. Our results suggest that these correlations reflect selection in response to NEAT domain-containing bacterial receptors like IsdB with conserved globin binding sites. A well-established body of literature has shown that other selective pressures play an important role in patterns of hemoglobin polymorphism and divergence across vertebrates, including adaptation to high altitude and malaria resistance (34, 38). It is therefore possible that signatures of selection detected in our study have been driven by pressures other than nutritional immunity. Nonetheless, our empirical results demonstrate that variation in hemoglobins at discrete sites has important functional consequences for bacterial iron acquisition.

Previous studies have illustrated how mutations in hemoglobin coding or regulatory regions can have highly deleterious effects on heme binding, oxygen affinity, and protein stability (43, 44). In addition to aforementioned sickle-cell alleles, dozens of hemoglobin mutations in humans have been reported that contribute to genetic disease, including anemia and thalassemia (45). Thus, despite identifying particular sites that are highly divergent among primates, much of the globin gene content is constrained due to purifying selection. In future work it would be useful to determine how variation among primate globins impacts other endogenous biochemical functions, such as heme binding and oxygen affinity. Such insights could improve our fundamental understanding of hemoglobin biology and the mechanisms underlying human hemoglobinopathies.

In conclusion, this work illustrates how rapid, site-specific hemoglobin variation restricts heme acquisition by the prominent human pathogen *S. aureus*. These findings provide a fundamental new perspective on vertebrate globin evolution, highlighting nutritional immunity as a selective pressure that could strongly impact divergence and natural selection. Future studies will assist in illuminating how these combinations of adaptive mutations contribute to hemoglobin function and host physiology. Understanding the genetic and molecular determinants of bacterial pathogenicity is critical for developing new antimicrobial treatment strategies, particularly as major pathogens like *S. aureus* continue to develop resistance to existing antibiotics. Combining comparative genetics with molecular experimentation in turn provides not only a historical perspective of host-microbe evolutionary conflict but also mechanistic insights on modern human infectious disease.

## ACKNOWLEDGEMENTS

We thank members of the Skaar and Barber laboratories for critical evaluation of this manuscript. We acknowledge the gift of pHUG21 from Douglas Henderson (University of Texas of the Permian Basin) and pHb0.0 from John Olson (Rice University).

## AUTHOR CONTRIBUTIONS

Conceptualization, J.E.C, E.P.S, M.F.B; Investigation, J.E.C., H.B.B., A.J.F, M.F.B; Writing-original draft, J.E.C, E.P.S, M.F.B.; Writing-reviewing and editing, J.E.C., H.B.B., A.J.F., E.P.S., M.F.B; Visualization, J.E.C, M.F.B; Supervision, E.P.S., M.F.B., Funding acquisition, E.P.S., M.F.B.

## MATERIALS AND METHODS

### Bacterial strains

For *E. coli* strains, LB agar and broth (Fisher, Hampton, NH) were routinely used and grown at 37°C. For selection of pHUG21, 12.5 μg/ml of carbenicillin (Fisher) was used; for selection of pHb0.0, 5 μg/ml of tetracycline hydrochloride (Alfa Aesar, Havermill, MA) was used, and for selection of pOS1 P_*lgt*_, 50 μg/ml of carbenicillin was used. *Staphylococcus* strains were grown at 37°C using tryptic soy agar and broth (Fisher), except when noted throughout. For selection of pOS1 P_*lgt*_ 10 μg/ml chloramphenicol (Fisher) was used. Strains were streaked to agar from stocks stored at −80°C two days prior to each experiment.

### Hemoglobin cloning and genetic manipulation

We compiled a subset of α- and β-globin sequences from GenBank, as well as cloned α-globin orthologs from cDNA derived from primate cell lines and β-globin orthologs from primate genomic DNA. Species obtained from GenBank: olive baboon, bonobo, white-headed capuchin, chimpanzee, Angolan colobus, northern white-cheeked gibbon, green monkey, human, crab-eating macaque, common marmoset, Sumatran orangutan, rhesus macaque, golden snub-nosed monkey, and squirrel monkey. The α-globin orthologs cloned from cDNA: African green monkey, black-and-white colobus, white-handed gibbon, Western lowland gorilla, Francois’ leaf monkey, black crested mangabey, white-faced marmoset, Nancy Ma’s night monkey, patas monkey, proboscis monkey, Allen’s swamp monkey, talapoin, and Wolf’s guenon. The β-globin orthologs cloned from genomic DNA: crested macaque, drill, Bolivian red howler monkey, pigtailed macaque, black-crested mangabey, Nancy Ma’s night monkey, patas monkey, white-faced saki, island siamang, black snub-nosed monkey, Allen’s swamp monkey, talapoin, Spix’s saddle-back tamarin, dusky titi, Wolf’s guenon, and common woolly monkey. Primate cell lines were purchased from the Coriell Institute for Medical Research (Camden, NJ).

Primate hemoglobin cDNA was cloned into pHb0.0 using Gibson assembly (New England Biolabs [NEB], Ipswich, MA). In general, each α- and β-globin gene cDNA was amplified from template (above) using Phusion 2X Master Mix (Thermo, Waltham, MA) with primers that also had homology to pHb0.0. Because of cDNA sequence homology, some primers were used for multiple species. pHb0.0-human was digested with PacI (NEB) and HindIII-HF (NEB) and the double-digested vector was isolated by gel purification (Qiagen, Germantown, MD). PCR products were assembled with digested pHb0.0, transformed to DH5α, re-isolated by mini-prep (Thermo) and were confirmed by sequencing (GeneWiz, South Plainfield, NJ) with pHb0.0_for/pHb0.0_rev. Globin cDNA was amplified for assembly as follows: white-cheeked gibbon α-globin-primers AF327/328, white-cheeked gibbon β-globin-primers AF329/330, baboon α-globin-primers AF331/332, baboon β-globin-primers AF329/330, talapoin α-globin-primers AF327/328, talapoin β-globin-primers AF329/330, marmoset α-globin-primers AF333/334, and marmoset β-globin-primers AF329/335.

Chimeric hemoglobins were prepared by subcloning the α-globins. To enable digestion with XbaI (NEB) (which is sensitive to *dam* methylation) pHb0.0-human and pHb0.0-baboon were transformed and re-isolated from *E. coli* K1077 (*dam^−^ dcm^−^*). The α-globin from each plasmid was excised by digestion with XbaI (NEB) and PacI (NEB) and the α-globin and double-digested pHb0.0 containing β-globin were separately isolated by gel purification (Qiagen). Human α-globin was ligated into pHb0.0 containing baboon β-globin and baboon α-globin was ligated (T4 ligase; NEB) into pHb0.0 containing human β-globin. Chimeras were confirmed by sequencing (GeneWiz) with pHb0.0_for/pHb0.0_rev.

pHb0.0-human was mutagenized using QuikChange Site-Directed Mutagenesis Kit (Agilent, Santa Clara, CA) to create changes in the α-gene: T8K (primers AF289/290), T8K;N9H (primers JC112/113; using pHb0.0-human αT8K as template), and A5D;T8K;N9H (primers JC114/115; using pHb0.0-human αT8K;N9H as template). Changes in the human β-gene were as follows: S9N (primers AF303/304), S9A (primers AF309/310) A76N (primers AF313/314), and A76T (primers AF311/312). Residues in pHb0.0-baboon α-gene were mutagenized: K8T (primers JC194/195), K8T;H9N simultaneously (primers JC116/117), and D5A (primers JC118/119; using pHb0.0-baboon αK8T;H9N as template).

### Phylogenetic analyses

Hemoglobin DNA sequence alignments were performed using MUSCLE. Input phylogenies were based upon supported species relationships (46) as well as maximum-likelihood gene phylogenies generated using PhyML with SPR topology search and 1000 bootstraps for branch support (47). Substitution models were selected based on the ProtTest algorithm (48). Tests for positive selection were performed using codeml from the PAML software package with the F3×4 codon frequency model. Likelihood ratio tests (LRTs) were performed by comparing pairs of site-specific models (NS sites): M1 (neutral) with M2 (selection), M7 (neutral, beta distribution of *d*N/*d*S<1) with M8 (selection, beta distribution, *d*N/*d*S>1 allowed). Additional tests which also account for synonymous rate variation and recombination, including FUBAR, FEL, and MEME, were performed using the HyPhy software package via the Datamonkey server (49, 50). Sites under positive selection were mapped onto three-dimensional molecular structures using Chimera (51) (http://www.cgl.ucsf.edu/chimera/).

The staphylococcal DNA gyrase gene tree was generated using PhyML with 1000 bootstraps as above. *M. caseolyticus* DNA gyrase was included as an outgroup. The similarity of IsdB in *S. argenteus* and *S. schweitzeri* relative to *S. aureus* is shown at right.

### Recombinant purification of hemoglobin

Hemoglobin expression strains [BL21(DE3) pHUG21 pHb0.0] were streaked to LB agar containing 12.5 μg/ml carbenicillin and 5 μg/ml tetracycline hydrochloride. Single colonies were inoculated into 5 ml of LB broth supplemented with 12.5 ug/ml carbenicillin and 5 ug/ml tetracycline hydrochloride and grown for 14 h at 37°C with shaking. This culture was used to inoculated 1:500 into 1.5 L of LB with 12.5 ug/ml carbenicillin, 5 ug/ml tetracycline hydrochloride, 100 μM hemin (prepared fresh at 10 mM in 0.1 M NaOH; Sigma St. Louis, MO), and 50 μg/ml of the iron chelator ethylenediamine-di(o-hydroxyphenylacetic acid (EDDHA [LGC Standards, Teddington, UK; solid added directly to medium) in a 2.8 L Fernbach flask. Cultures were grown at 37°C until OD_600_ reached 0.6-0.8. The expression of hemoglobin was induced with 40 μg/ml IPTG (RPI, Mount Prospect, IL). After 16 h post-induction, cells were collected by centrifugation. The cell pellet was resuspended in PBS containing 10 mM imidazole (Fisher), 5 mM MgCl_2_ (Sigma), 1 Roche Protease inhibitor table (Fisher), and a few mg of lysozyme (Thermo) and deoxyribonuclease from bovine pancreas (Sigma). The cell pellet resuspended with rocking for 20 min following incubation on ice. Cells were lysed using an Emulsiflex (Avestin, Ottawa, CA) then cell lysate was clarified by ultracentrifugation (60 min at 17,000 g). Cell lysate was applied to a Ni-NTA resin (Qiagen), to which hemoglobin binds, washed with PBS with10 mM imidazole. Hemoglobin was eluted with PBS with 500 mM imidazole. The hemoglobin-containing eluate was dialyzed twice sequentially in PBS at 4°C. Purified hemoglobin was filter sterilized with a 0.45 micron filter and stored in aliquots in liquid nitrogen. Hemoglobin concentration was measured with Drabkin’s reagent (Sigma) using human hemoglobin as a standard.

### Whole cell hemoglobin binding assay

*S. aureus* strains were streaked on tryptic soy agar (containing 10 μg/ml chloramphenicol for strains carrying plasmids) and grown at 37°C for 24 h. Single colonies were used to inoculate 3 ml of RPMI containing 1%cas-amino acids and 0.5 mM 2,2’-dipyridyl (Acros/Fisher) to induce expression of chromosomal *isdB*) or 10 μg/ml chloramphenicol (for strains carrying plasmids with constitutive *isdB* expression). After 14-16 h of growth at 37°C with shaking, 2 OD_600_ units were collected by centrifugation in a 1.5 ml Eppendorf tube. The cell pellet was resuspended with 1 ml PBS or PBS containing recombinant hemoglobin. 10 μg/ml (chromosomal IsdB) or 2.5 μg/ml (plasmid-borne IsdB) of hemoglobin was used. The cells were incubated with hemoglobin or PBS for 30 min at 37°C with shaking, then cells were collected by centrifugation at 4°C. Cells were washed thrice with 1 ml ice-cold PBS, centrifuging at 4°C. After the final wash, the cells were resuspended in 30 μl 0.5 M Tris pH 8.0 (Fisher) containing 4% SDS (Fisher) and heated at 90°C for 5 min to remove surface bound proteins. Cells were collected by centrifugation, and eluate was added to 6X loading buffer and heated at 90°C for 5 min. Samples were subjected to 12 or 17.5% SDS-PAGE and silver stained (GE, Boston, MA). Quantification was performed by densitometric analysis with Image J (NIH). Because of variation in stain intensity across gels, all comparisons were made within the same gel, and relative density was calculated for each biological replicate within the same gel; the comparison was either to human hemoglobin or wildtype IsdB, depending on assay. Additionally, PBS only samples and *S. aureus* Δ*isdB::erm* were used to verify that hemoglobin binding in this assay is IsdB dependent as previously observed [Supplemental Figure 1] (11, 13).

### Growth with hemoglobin as sole iron source

*S. aureus* Newman WT was streaked to tryptic soy agar and allowed to grow for 24 h at 37°C. A few colonies were used to inoculate 5 ml of RPMI (Corning, Corning, NY) supplemented with 1% casamino acids (Fisher) and 0.5 mM of EDDHA (prepared fresh in ethanol). After growth to stationary phase at 37°C with shaking, approximately 16 h, 4 μl of culture was inoculated into 196 μl of medium in a 96 well plate and OD_600_ at 37°C with shaking was monitored over time using a BioTek plate reader. Medium was RPMI containing 1% cas-amino acids that had been stripped of cations with Chelex 100 (Sigma) according to manufacturer’s instructions, filter sterilized, and supplemented with 25 μM ZnCl_2_, 25 μM MnCl_2_, 100 μM CaCl_2_, and 1 mM MgCl_2_ (all from Fisher) to restore non-iron cations, 1.5 mM EDDHA to chelate any remaining free iron, and 2.5 μg/ml of recombinant purified hemoglobin as the sole iron source. Growth using hemoglobin as a sole iron source is IsdB dependent (11, 13, 22).

### *Cloning of* isdB

The full length coding sequences of IsdB were amplified from genomic DNA using Phusion 2X High-Fidelity Master Mix (Thermo); cells were treated with 20 μg of lysostaphin (AMBI Products, Lawrence, NY) and DNA wasisolated with Wizard genomic DNA extraction kit (Promega, Madison, WI). *S. aureus isdB* (*NWMN_1040*) was amplified using primers JC343/344, *S. schweitzeri isdB* (*ERS140239_01018*) using primers JC218/219, and *S. argenteus isdB* (*SAMSHR1132_09750*) using primers JC216/217. Each primer pair included homology to pOS1 P_*lgt*_ digested with NdeI and BamHI-HF (NEB), and PCR products were ligated to pOS1 P_*lgt*_ with Hi-Fi assembly (NEB), transformed into *E. coli* DH5α and re-isolated by miniprep (Thermo). All plasmids were sequence confirmed by Sanger sequencing (Genewiz). Plasmids were transformed into RN4220 by electroporation, re-isolated, and transformed into *S. aureus* Newman Δ*isdB::erm* by electroporation.

### S. aureus *IsdB site-directed mutagenesis*

pOS1 P*_lgt_isdB_aureus_* was subjected to site-directed mutagenesis by PCR with Q5 Site Directed Mutagenesis (NEB). Primer pairs with desired mutation were used to create Q162R (primers JC315/316), S170T (primers 317/318) and Q162R;S170T (primers JC319/320). PCR products were transformed to DH5α. Plasmids were isolated and subjected to Sanger sequencing with primers JC228/229 (Genewiz) to identify successful incorporation of the desired mutation. Plasmids were transformed into RN4220 by electroporation, re-isolated, and transformed into *S. aureus* Newman Δ*isdB::erm* by electroporation.

## QUANTIFICATION AND STATISTICAL ANALYSIS

Specific statistical details for each experiment can be found in the corresponding figure legend. Data analysis and statistical tests were performed in Prism 6 (Graphpad).

**Table S1.** Strains and plasmids

| Bacterial Strain | Source |
| --- | --- |
| *Escherichia coli* DH5α | Laboratory stock, Thermo Fisher |
| *Escherichia coli* BL21(DE3) pHUG21 | Douglas Henderson (University of Texas of the Permian Basin); (52) |
| *Staphylococcus aureus* RN4220 | (53) |
| *S. aureus* Newman | Laboratory stock; (54) |
| *S. aureus* Newman $\Delta isdB::erm$ | (7) |
| *S. argenteus* MSHR1132 | DSMZ |
| *S. schweitzeri* FSA084 | DSMZ |
| Plasmid | Source |
| pHb0.0-human | John Olson (Rice University); (52) |
| pHb0.0-white-cheeked gibbon | This paper |
| pHb0.0-baboon | This paper |
| pHb0.0-talapoin | This paper |
| pHb0.0-marmoset | This paper |
| pHb0.0-$\alpha_{\text{human}}\beta_{\text{baboon}}$ | This paper |
| pHb0.0-$\alpha_{\text{baboon}}\beta_{\text{human}}$ | This paper |
| pHb0.0-human αT8K | This paper |
| pHb0.0-human αA5D;T8K;N9H | This paper |
| pHb0.0-baboon αK8T | This paper |
| pHb0.0-baboon αD5A;K8T;H9N | This paper |
| pHb0.0-baboon βS9N | This paper |
| pHb0.0-baboon βS9A | This paper |
| pHb0.0-baboon βA76N | This paper |
| pHb0.0-baboon βA76T | This paper |
| pOS1 P<sub>Igt</sub> | (55) |
| pOS1 P<sub>Igt</sub>*isdB*<sub>aureus</sub> | This paper |
| pOS1 P<sub>Igt</sub>*isdB*<sub>schweitzeri</sub> | This paper |
| pOS1 P<sub>Igt</sub>*isdB*<sub>argenteus</sub> | This paper |
| pOS1 P<sub>Igt</sub>*isdB*<sub>aureus</sub> Q162R | This paper |
| pOS1 P<sub>Igt</sub>*isdB*<sub>aureus</sub> S170T | This paper |
| pOS1 P<sub>Igt</sub>*isdB*<sub>aureus</sub> Q162R;S170T | This paper |

**Table S2.**
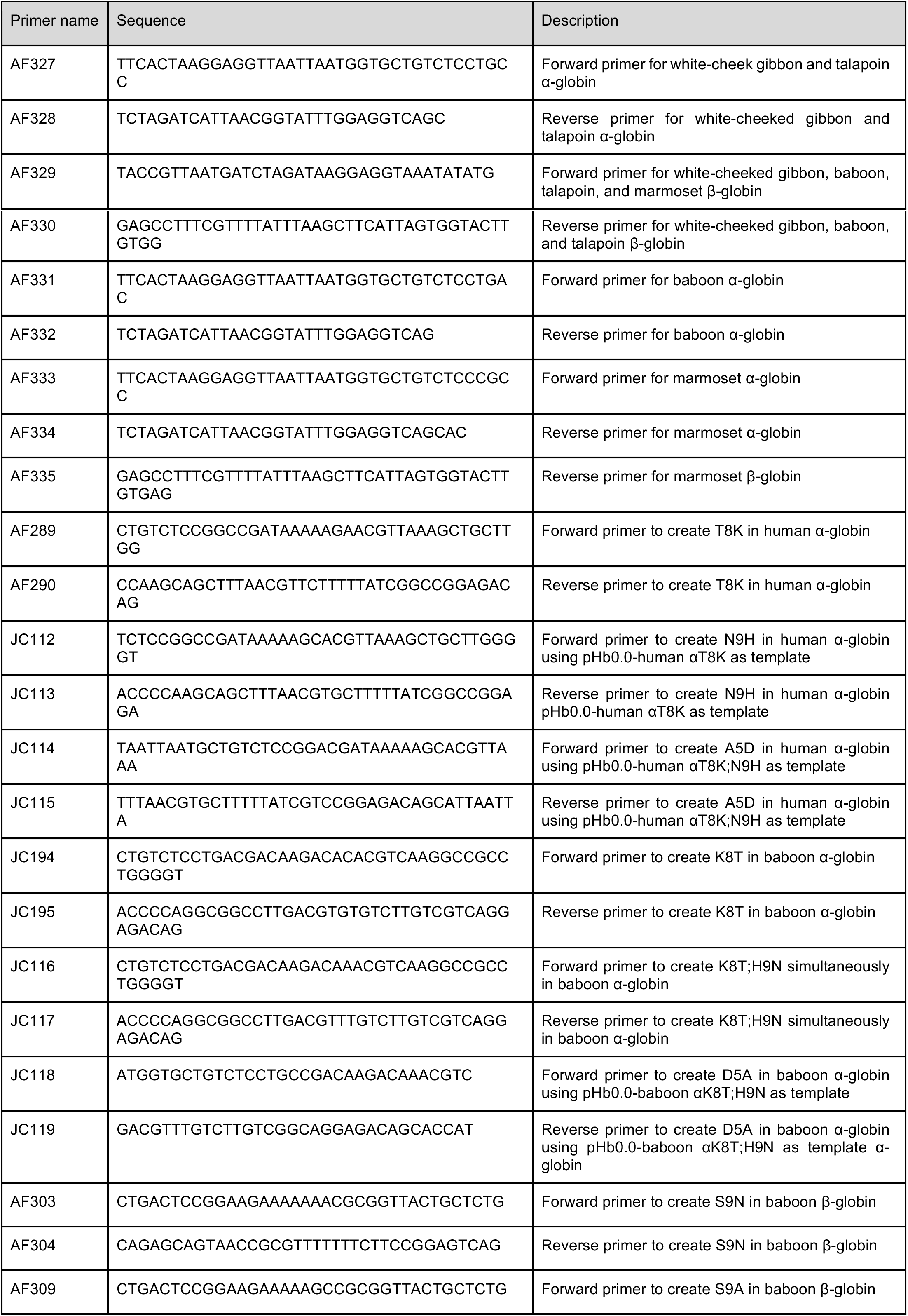
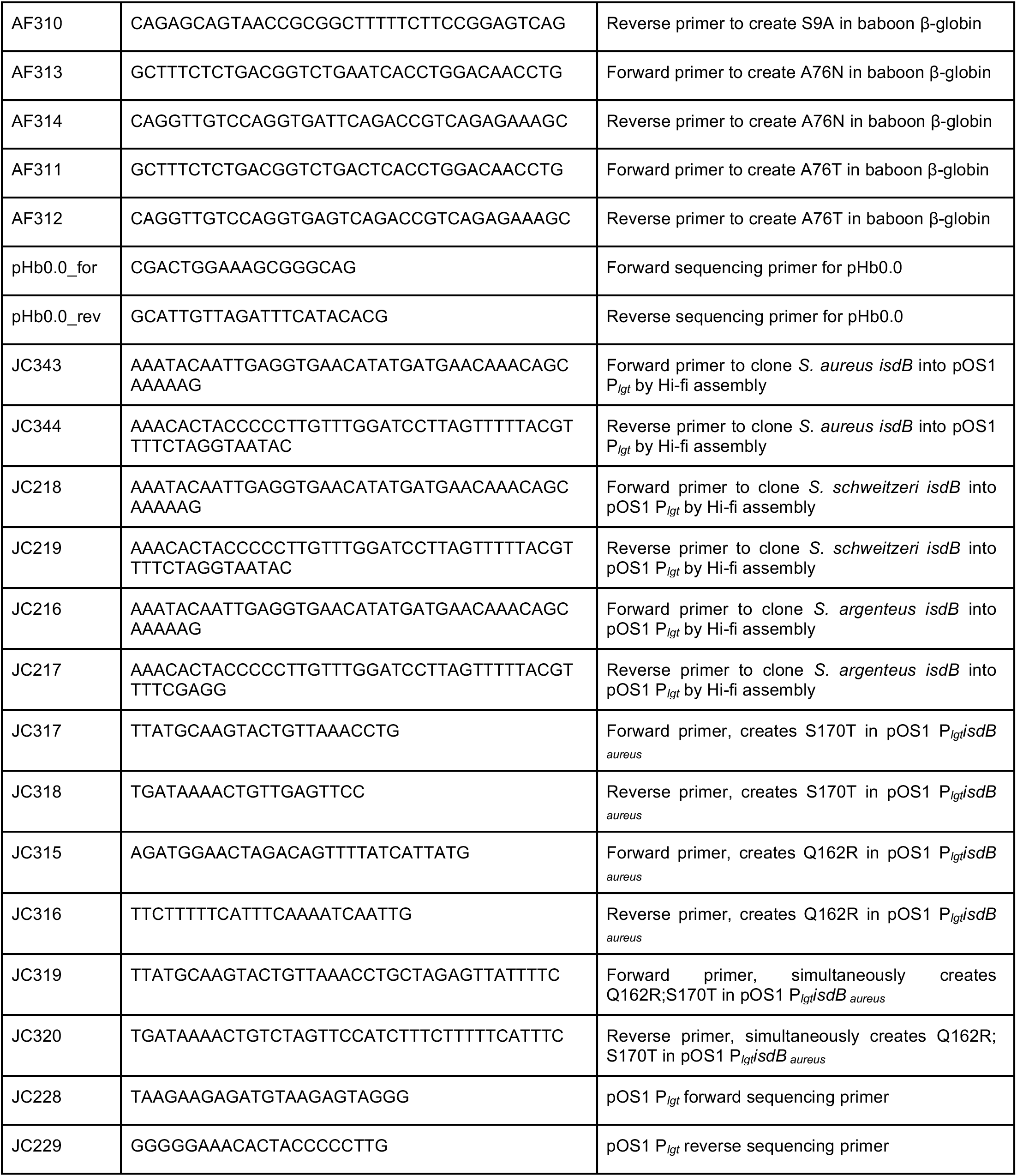
Oligonucleotides used in this study

## SUPPLEMENTAL INFORMATION

**Supplemental Figure 1.**
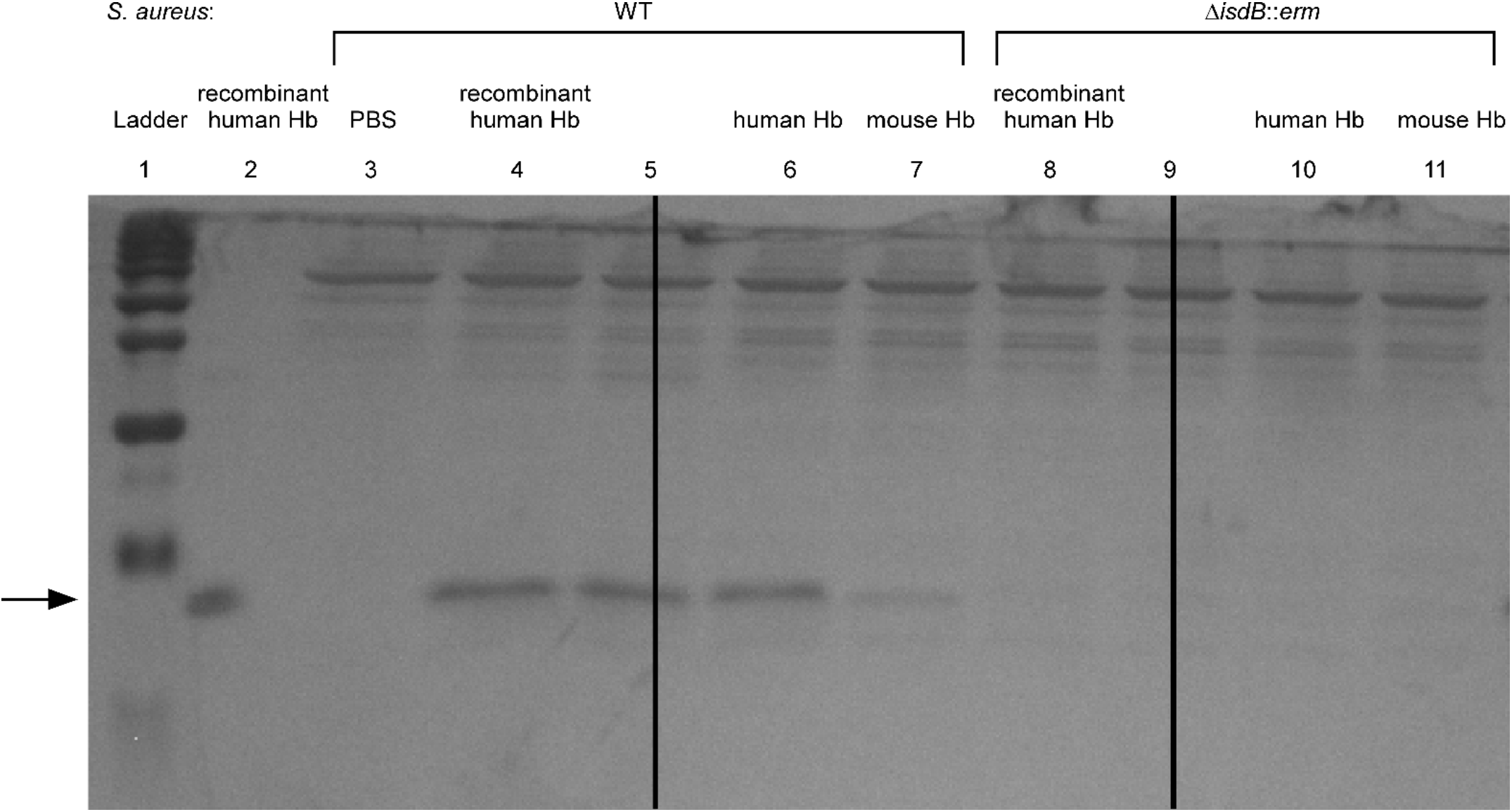
The whole cell hemoglobin (Hb) binding assay allows for IsdB-dependent, species-specific quantification of bound recombinant Hb. A representative silver-stained SDS-PAGE gel used to assess binding of hemoglobin to the surface of *S. aureus*. 100 ng of purified recombinant Hb was loaded (lane 2) to demonstrate apparent molecular weight in this gel. The apparent hemoglobin band is specific to Hb, as it is not visible when *S. aureus* is incubated with PBS alone (lane 3). *S. aureus* binds human hemoglobin purified recombinantly from *E. coli* (lane 4) as well as Hb purified from blood (lane 6). Binding of Hb purified from mouse blood (lane 7) is significantly diminished, as first reported in (13) Binding of Hb is dependent on IsdB, as demonstrated in lanes 8,10,11. An approximately equivalent amount of *S. aureus* was used, as demonstrated by equal loading of non-Hb specific bands across the top of the gel. Samples in lanes 5 and 9 are not discussed here.

## REFERENCES

1. Palmer LD, Skaar EP. Transition Metals and Virulence in Bacteria. Annu Rev Genet. 2016;50:67–91.

2. Weinberg ED. Nutritional immunity: host’s attempt to withold iron from microbial invaders. JAMA. 1975;231(1):39–41.

3. Schade AL, Caroline L. Raw Hen Egg White and the Role of Iron in Growth Inhibition of *Shigella dysenteriae, Staphylococcus aureus, Escherichia coli and Saccharomyces cerevisiae*. Science. 1944;100(2584):14–5.

4. Cassat JE, Skaar EP. Iron in infection and immunity. Cell Host Microbe. 2013;13(5):509–19.

5. Choby JE, Skaar EP. Heme synthesis and acquisition in bacterial pathogens. J Mol Biol. 2016;428(17):3408–28.

6. Klevens RM, Morrison MA, Nadle J, Petit S, Gershman K, Ray S, et al. Invasive methicillin-resistant *Staphylococcus aureus* infections in the United States. JAMA. 2007;298(15):1763–71.

7. Mazmanian SK, Skaar EP, Gaspar AH, Humayun M, Gornicki P, Jelenska J, et al. Passage of heme-iron across the envelope of *Staphylococcus aureus*. Science. 2003;299:906–9.

8. Spaan AN, Reyes-Robles T, Badiou C, Cochet S, Boguslawski KM, Yoong P, et al. *Staphylococcus aureus* targets the Duffy Antigen Receptor for Chemokines (DARC) to lyse erythrocytes. Cell Host Microbe. 2015;18(3):363–70.

9. Torres VJ, Pishchany G, Humayun M, Schneewind O, Skaar EP. *Staphylococcus aureus* IsdB is a hemoglobin receptor required for heme iron utilization. J Bacteriol. 2006;188(24):8421–9.

10. Dryla A, Gelbmann D, Von Gabain A, Nagy E. Identification of a novel iron regulated staphylococcal surface protein with haptoglobin-haemoglobin binding activity. Molecular Microbiology. 2003;49(1):37–53.

11. Pishchany G, Sheldon JR, Dickson CF, Alam MT, Read TD, Gell DA, et al. IsdB-dependent hemoglobin binding is required for acquisition of heme by *Staphylococcus aureus*. J Infect Dis. 2014;209(11):1764–72.

12. Cheng AG, Kim HK, Burts ML, Krausz T, Schneewind O, Missiakas DM. Genetic requirements for *Staphylococcus aureus* abscess formation and persistence in host tissues. FASEB J. 2009;23(10):3393–404.

13. Pishchany G, McCoy AL, Torres VJ, Krause JC, Crowe JE, Jr., Fabry ME, et al. Specificity for human hemoglobin enhances *Staphylococcus aureus* infection. Cell Host Microbe. 2010;8(6):544–50.

14. Malachowa N, Whitney AR, Kobayashi SD, Sturdevant DE, Kennedy AD, Braughton KR, et al. Global changes in *Staphylococcus aureus* gene expression in human blood. PLoS One. 2011;6(4):e18617.

15. Fowler VG, Jr., Proctor RA. Where does a *Staphylococcus aureus* vaccine stand? Clin Microbiol Infect. 2014;20 Suppl 5:66–75.

16. Muryoi N, Tiedemann MT, Pluym M, Cheung J, Heinrichs DE, Stillman MJ. Demonstration of the iron-regulated surface determinant (Isd) heme transfer pathway in *Staphylococcus aureus*. J Biol Chem. 2008;283(42):28125–36.

17. Skaar EP, Gaspar AH, Schneewind O. IsdG and IsdI, heme-degrading enzymes in the cytoplasm of *Staphylococcus aureus*. J Biol Chem. 2004;279(1):436–43.

18. Grigg JC, Vermeiren CL, Heinrichs DE, Murphy ME. Heme coordination by *Staphylococcus aureus* IsdE. J Biol Chem. 2007;282(39):28815–22.

19. Zhu H, Xie G, Liu M, Olson JS, Fabian M, Dooley DM, et al. Pathway for heme uptake from human methemoglobin by the iron-regulated surface determinants system of *Staphylococcus aureus*. J Biol Chem. 2008;283(26):18450–60.

20. Reniere ML, Ukpabi GN, Harry SR, Stec DF, Krull R, Wright DW, et al. The IsdG-family of haem oxygenases degrades haem to a novel chromophore. Mol Microbiol. 2010;75(6):1529–38.

21. Skaar EP, Humayun M, Bae T, DeBord KL, Schneewind O. Iron-source preference of *Staphylococcus aureus* infections. Science. 2004;305:1626–8.

22. Bowden CFM, Chan ACK, Li EJW, Arrieta AL, Eltis LD, Murphy MEP. Structure-function analyses reveal key features in *Staphylococcus aureus* IsdB-associated unfolding of the heme-binding pocket of human hemoglobin. J Biol Chem. 2018;293(1):177–90.

23. Daugherty MD, Malik HS. Rules of engagement: molecular insights from host-virus arms races. Annu Rev Genet. 2012;46:677–700.

24. Hughes AL, Nei M. Pattern of nucleotide substitution at major histocompatibility complex class I loci reveals overdominant selection. Nature. 1988;335(6186):167–70.

25. Hamilton WD, Axelrod R, Tanese R. Sexual reproduction as an adaptation to resist parasites (a review). Proc Natl Acad Sci U S A. 1990;87(9):3566–73.

26. Yang Z, Bielawski JP. Statistical methods for detecting molecular adaptation. Trends in ecology & evolution. 2000;15(12):496–503.

27. Nei M, Gojobori T. Simple methods for estimating the numbers of synonymous and nonsynonymous nucleotide substitutions. Molecular biology and evolution. 1986;3(5):418–26.

28. Demogines A, Farzan M, Sawyer SL. Evidence for ACE2-utilizing coronaviruses (CoVs) related to severe acute respiratory syndrome CoV in bats. J Virol. 2012;86(11):6350–3.

29. Sawyer SL, Wu LI, Emerman M, Malik HS. Positive selection of primate TRIM5alpha identifies a critical species-specific retroviral restriction domain. Proc Natl Acad Sci U S A. 2005;102(8):2832–7.

30. Elde NC, Child SJ, Geballe AP, Malik HS. Protein kinase R reveals an evolutionary model for defeating viral mimicry. Nature. 2009;457(7228):485–9.

31. Enard D, Cai L, Gwennap C, Petrov DA. Viruses are a dominant driver of protein adaptation in mammals. eLife. 2016;5.

32. Barber MF, Elde NC. Nutritional immunity. Escape from bacterial iron piracy through rapid evolution of transferrin. Science. 2014;346(6215):1362–6.

33. Barber MF, Kronenberg Z, Yandell M, Elde NC. Antimicrobial Functions of Lactoferrin Promote Genetic Conflicts in Ancient Primates and Modern Humans. PLoS Genet. 2016;12(5):e1006063.

34. Storz JF, Moriyama H. Mechanisms of hemoglobin adaptation to high altitude hypoxia. High Alt Med Biol. 2008;9(2):148–57.

35. Galen SC, Natarajan C, Moriyama H, Weber RE, Fago A, Benham PM, et al. Contribution of a mutational hot spot to hemoglobin adaptation in high-altitude Andean house wrens. Proc Natl Acad Sci U S A. 2015;112(45):13958–63.

36. Natarajan C, Hoffmann FG, Weber RE, Fago A, Witt CC, Storz JF. Predictable convergence in hemoglobin function has unpredictable molecular underpinnings. Science. 2016;354(6310):336–9.

37. Storz JF, Sabatino SJ, Hoffmann FG, Gering EJ, Moriyama H, Ferrand N, et al. The molecular basis of high-altitude adaptation in deer mice. PLoS Genet. 2007;3(3):e45.

38. Ferreira A, Marguti I, Bechmann I, Jeney V, Chora A, Palha NR, et al. Sickle hemoglobin confers tolerance to *Plasmodium* infection. Cell. 2011;145(3):398–409.

39. Kloos WE. Natural populations of the genus *Staphylococcus*. Annu Rev Microbiol. 1980;34:559–92.

40. Wright JS, 3rd, Traber KE, Corrigan R, Benson SA, Musser JM, Novick RP. The agr radiation: an early event in the evolution of staphylococci. J Bacteriol. 2005;187(16):5585–94.

41. van Belkum A, Melles DC, Nouwen J, van Leeuwen WB, van Wamel W, Vos MC, et al. Co- evolutionary aspects of human colonisation and infection by *Staphylococcus aureus*. Infect Genet Evol. 2009;9(1):32–47.

42. Massey RC, Horsburgh MJ, Lina G, Hook M, Recker M. The evolution and maintenance of virulence in *Staphylococcus aureus:* a role for host-to-host transmission? Nat Rev Microbiol. 2006;4(12):953–8.

43. Weatherall D, Akinyanju O, Fucharoen S, Olivieri N, Musgrove P. Inherited Disorders of Hemoglobin. In: Jamison DT, Breman JG, Measham AR, Alleyne G, Claeson M, Evans DB, et al., editors. Disease Control Priorities in Developing Countries. 2nd ed. Washington (DC): Oxford University Press and The World Bank; 2006. p. 663–80.

44. Stamatoyannopoulos G. The molecular basis of hemoglobin disease. Annu Rev Genet. 1972;6:47–70.

45. Thom CS, Dickson CF, Gell DA, Weiss MJ. Hemoglobin variants: biochemical properties and clinical correlates. Cold Spring Harb Perspect Med. 2013;3(3):a011858.

46. Perelman P, Johnson WE, Roos C, Seuanez HN, Horvath JE, Moreira MA, et al. A molecular phylogeny of living primates. PLoS Genet. 2011;7(3):e1001342.

47. Guindon S, Dufayard JF, Lefort V, Anisimova M, Hordijk W, Gascuel O. New algorithms and methods to estimate maximum-likelihood phylogenies: assessing the performance of PhyML 3.0. Systematic biology. 2010;59(3):307–21.

48. Abascal F, Zardoya R, Posada D. ProtTest: selection of best-fit models of protein evolution. Bioinformatics. 2005;21 (9):2104–5.

49. Delport W, Poon AF, Frost SD, Kosakovsky Pond SL. Datamonkey 2010: a suite of phylogenetic analysis tools for evolutionary biology. Bioinformatics. 2010;26(19):2455–7.

50. Pond SL, Frost SD, Muse SV. HyPhy: hypothesis testing using phylogenies. Bioinformatics. 2005;21(5):676–9.

51. Pettersen EF, Goddard TD, Huang CC, Couch GS, Greenblatt DM, Meng EC, et al. UCSF Chimera--a visualization system for exploratory research and analysis. Journal of computational chemistry. 2004;25(13):1605–12.

52. Villarreal DM, Phillips CL, Kelley AM, Villarreal S, Villaloboz A, Hernandez P, et al. Enhancement of recombinant hemoglobin production in *Escherichia coli* BL21(DE3) containing the *Plesiomonas shigelloides* heme transport system. Applied and environmental microbiology. 2008;74(18):5854–6.

53. Kreiswirth BN, Lofdahl S, Betley MJ, O’Reilly M, Schlievert PM, Bergdoll MS, et al. The toxic shock syndrome exotoxin structural gene is not detectably transmitted by a prophage. Nature. 1983;305(5936):709–12.

54. Duthie EL, Lisa L. Staphylococcal coagulase: mode of action and antigenicity. J Gen Microbiol. 1952;6:95–107.

55. Bubeck Wardenburg J, Williams WA, Missiakas D. Host defenses against *Staphylococcus aureus* infection require recognition of bacterial lipoproteins. Proc Natl Acad Sci U S A. 2006;103(37):13831–6.

